# Cohesin regulates promoter-proximal pausing of RNA Polymerase II by limiting recruitment of super elongation complex

**DOI:** 10.1101/2024.03.15.584908

**Authors:** Shoin Tei, Toyonori Sakata, Atsunori Yoshimura, Toyoaki Natsume, Masato T Kanemaki, Masashige Bando, Katsuhiko Shirahige

**Affiliations:** Laboratory of Genome Structure and Function, Institute for Quantitative Biosciences, The University of Tokyo, 1-1-1 Yayoi, Bunkyo-Ku, Tokyo, 113-0032, Japan; Karolinska Institutet, Department of Biosciences and Nutrition, Biomedicum, Quarter A6, 171 77, Stockholm, Sweden; Karolinska Institutet, Department of Cell and Molecular Biology, Biomedicum, Quarter A6, 171 77, Stockholm, Sweden; Department of Chromosome Science, National Institute of Genetics, Research Organization of Information and Systems (ROIS), Yata 1111, Mishima, Shizuoka 411-8540, Japan; Graduate Institute for Advanced Studies, SOKENDAI, Yata 1111, Mishima, Shizuoka 411-8540, Japan; Research Center for Genome & Medical Sciences, Tokyo Metropolitan Institute of Medical Science, Tokyo, Japan; Department of Biological Science, The University of Tokyo, Bunkyo-ku, Tokyo 113-0033, Japan

## Abstract

Cohesin is a ring-shaped complex, responsible for establishing sister chromatid cohesion and forming topologically associating domains (TADs) and chromatin loops. Loss-of-function mutations in cohesin subunits and its regulatory factors can cause Cornelia de Lange syndrome (CdLS). Because dysregulated gene expression was observed in CdLS, it has long been thought that cohesin plays a regulatory role in transcription. Here, we investigated the effect of acute cohesin depletion on transcription and observed that a small number of genes exhibited differential expression. Analysis of RNA polymerase II (Pol II) distribution revealed that the depletion reduced Pol II promoter binding and pausing simultaneously at the majority of genes. This implies that at most genes, the two decreases counterbalance each other, resulting in unchanged gene expression. Additionally, we find that cohesin loss increased promoter binding of super elongation complex (SEC), which mediates the release of Pol II from paused state. Moreover, the reduction in pausing caused by cohesin depletion was no longer observed when SEC was inhibited. These observations suggest that cohesin regulates Pol II pausing by restricting SEC recruitment to promoters. Together, our study demonstrates the involvement of cohesin in transcriptional regulation, particularly in Pol II pause and release.

## Introduction

Cohesin is a ring-shaped complex consisting of four core subunits, SMC1, SMC3, Rad21, and SA1/2. Nipbl/Mau2 complex as cohesin loader facilitates the binding of cohesin to DNA.^1^ Cohesin was initially identified for its essential role in sister chromatid cohesion.^2,3,4^ Cohesin holds sister chromatids together from S phase until anaphase by trapping DNA inside its ring for proper chromosome segregation. Chromatin conformation capture (3C)-based methods such as Hi-C and Micro-C have revealed that interphase chromosomes are organized into 3D structures at multiple levels through cis-interactions.^5,6,7,8^ Among the architectural features, cohesin actively contributes to the formation of topologically associated domains (TADs) and chromatin loops.^9,10,11,12,13^ TADs are large genomic domains characterized by intensive chromatin interactions within themselves and limited contacts with the surrounding regions.^5^ Enriched binding of cohesin, together with CTCF (CCCTC-binding factor), is often detected at the boundaries of TADs and the anchors of loops. Recent studies, based on simulation or depletion of cohesin regulatory factors, support a model in which the loop extrusion activity of cohesin is responsible for the formation of the TADs and loops.^11,14,15,10,16,17^ In this model, cohesin slides along chromosomes while DNA loops are pushed through the cohesin ring. Then, the movement is blocked by the encounter with CTCF. Previous studies using single-molecule imaging observed the extruding cohesin on tethered DNA molecules.^18,19^ On the other hand, cohesin mutants unable to exclude DNA have been observed to form chromatin loops *in vivo*.^20^ Thus, the involvement of the loop extrusion activity in the formation of cohesin-mediated structures is still under debate. Cohesin is thought to regulate gene expression by modulating the contact frequency between promoters and enhancers through TADs and promoter-enhancer loops.^21^ Perturbations of TAD boundaries can cause ectopic promoter-enhancer interaction and lead to up-regulated expression of the gene.^22,23,24^ However, only a small part of genes exhibits differential expression levels upon cohesin depletion.^9,25,12,13,26^ Thus, the contribution of cohesin-mediated promoter-enhancer interactions to gene expression remains unclear.

Loss-of-function mutations in cohesin subunits and its regulatory factors cause Cornelia de Lange syndrome (CdLS), a congenital disorder characterized by developmental defects.^27,28^ Cohesin-related CdLS causative genes include Nipbl, SMC1, SMC3, Rad21, and HDAC8, with Nipbl accounting for more than 60% of cases. We demonstrated that CdLS share phenotype and dysregulated genes with CHOPS, which is associated with gain-of-function mutations in AFF4 gene.^29^ AFF4 is a component of super elongation complex (SEC), which regulates transcriptional productive elongation.^30,31^ The resemblance between CdLS and CHOPS suggests that dysregulated transcription might be a common underlying mechanism. Moreover, we propose that both these mutated cohesin factors and SEC operate in the same transcriptional pathway. In higher eukaryotes, shortly after transcription initiation, RNA polymerase II (Pol II) temporally pauses just downstream of transcription start site (TSS) with the help of pausing factors, such as negative elongation factor (NELF) and DRB sensitivity inducing factor (DSIF).^32,33^ Cdk9, within SEC, phosphorylates the Ser2 of Pol II C-terminal domain (CTD) (Ser2P), facilitating the release of paused Pol II into productive elongation. Simultaneously, Cdk9 phosphorylates NELF, which dissociates NELF from the transcriptional machinery. Inhibition of SEC is reported to impair Pol II release from the paused state and slow elongation along genes.^31^ Based on the above, our hypothesis posits that cohesin also plays a regulatory role in Pol II pausing and elongation. To study this hypothesis, we acutely depleted cohesin by targeted degradation of Rad21 and analyzed the effect of the depletion on gene expression, the genomic distribution of Pol II, and factors involved in the pausing and release, along with 3D chromatin structures. At the majority of genes, we find cohesin promotes promoter binding of Pol II and mediates pausing simultaneously. Furthermore, cohesin restricts SEC recruitment to promoters. This study introduces the involvement of cohesin in transcriptional regulation, particularly in Pol II pause and release.

## Results

### Cohesin regulates the expression of a subset of genes while promoting Pol II binding at the majority of promoters

To study the transcriptional regulation by cohesin, we examined the effect of acute cohesin depletion on gene expression and distribution of RNA polymerase II (Pol II) along genes. We employed the auxin-degron system (AID) to rapidly degrade Rad21 without affecting cell cycle progression (Fig. S1a-S1c).^9^ Quantification of band intensity revealed that Rad21 levels were reduced to about 20% of that in control cells. For a reliable analysis of the transcriptome and intragenic dynamics of Pol II, it is necessary to eliminate the influence of closely spaced or partially overlapping genes. In addition, the length of the genes should be greater than a certain length (2 kb) and the gene must be clearly expressed. Thus, we selected 6797 highly expressed and well-separated genes based on the criteria (see methods). For better quantification, 5-ethynyl 2’-deoxyuridine (EU)-labeled nascent RNA sequencing (EU-seq) and chromatin immunoprecipitation sequencing (ChIP-seq) were performed with spike-in normalization control. To observe the direct effect of the depletion on the transcriptome, we conducted EU-seq, where EU was introduced into the culture media for the last 20 min of the auxin treatment to label nascent RNA. Gene expression analysis (FDR < 0.05) identified only 250 differentially expressed genes (85 up- and 165 down-regulated) upon auxin treatment (Figs. 1a, S1d and S1e). The remaining 6547 genes did not exhibit significant changes in expression, indicating that rapid cohesin depletion had a minor effect on the transcriptome as previously reported.^9,25^ In contrast, ChIP-seq of RPB1, the largest subunit of Pol II revealed that cohesin depletion decreased Pol II at the majority of gene promoters. Specifically, we observed a significant reduction at 4463 promoters, an increase at 301, and no changes at 2033 genes (Fig. 1b). In line with the overall reduction at promoters, even upDEGs exhibited a slight decrease in RPB1 occupancy, while downDEGs showed the largest reduction (Fig. 1c). Therefore, we observed that cohesin loss had a minimal effect on gene expression despite the reduction in Pol II binding at most promoters. To resolve this discrepancy, we compared the changes in RPB1 density between promoters and gene bodies by linear regression.

**Fig. 1:**
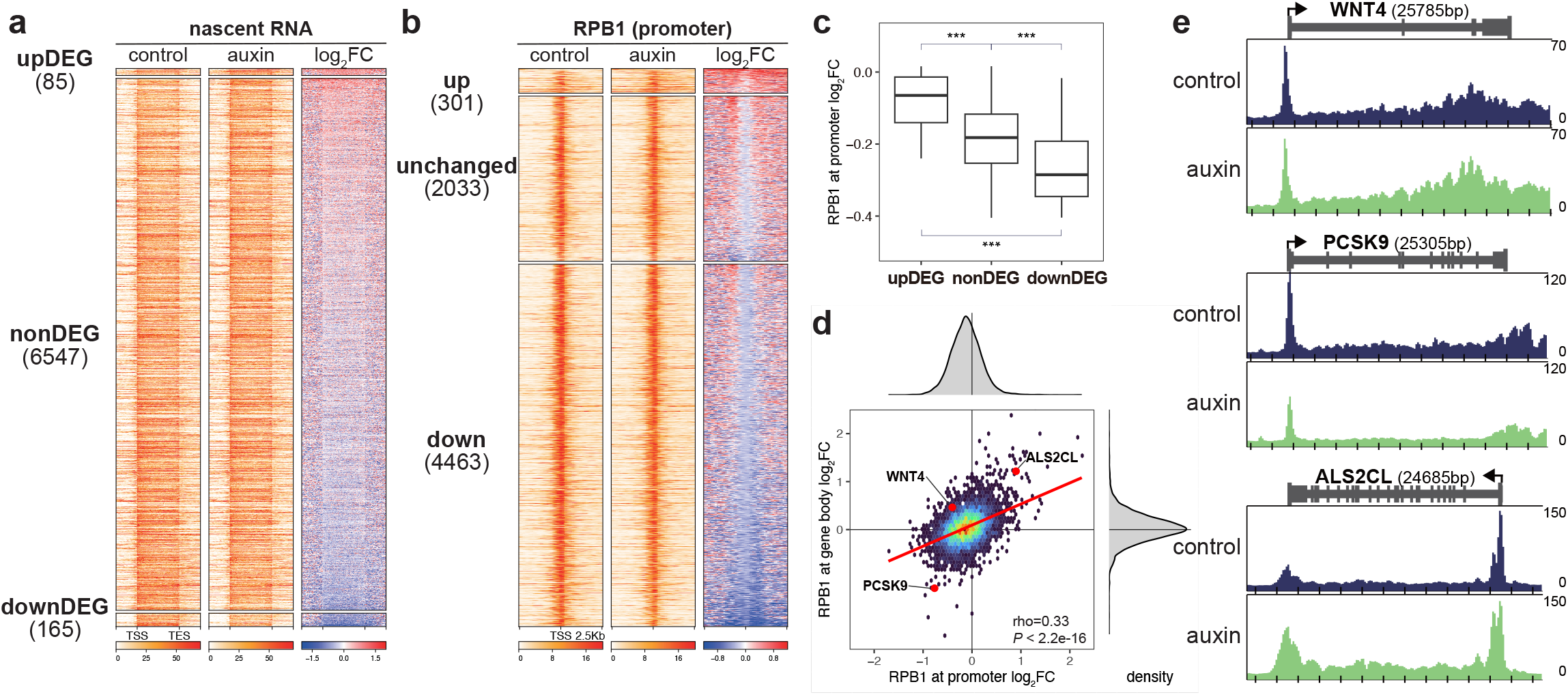
Cohesin regulates the expression of a subset of genes while promoting Pol II binding at the majority of promoters. **a** Heatmaps showing signals of nascent RNA in control and auxin-treated cells at upDEGs, nonDEGs, and downDEGs sorted by log2 fold change between control and auxin-treated cells. Genes analyzed in this study should have H3K4me3 peaks within 5kb of their TSSs, with RPKM of RPB1 ≥ 1, > 2kb in length, and >1kb away from the other genes. Differentially expressed genes (DEGs) were identified in comparison between control and auxin-treated cells using edgeR (FDR < 0.05). **b** Heatmaps showing signals of RPB1 in control and auxin-treated cells at promoters of genes with up-regulated, unchanged, and down-regulated RPB1 promoter occupancy upon auxin treatment, sorted by log2 fold change between control and auxin-treated cells. Genes were grouped into up-regulated, unchanged, and down-regulated using a binomial test comparing RPB1 promoter occupancy (-1 to 1kb) between control and auxin-treated cells (FDR < 0.05). FDR was calculated by adjusting P values for multiple testing using the Benjamini–Hochberg method. **c** Box plot showing the distribution of log2 fold change in RPB1 promoter occupancy upon auxin treatment at upDEGs, nonDEGs, and downDEGs. A multiple Welch t-test was performed between every pair of gene groups. ***P[<[0.001. P = 3.8e-13 (upDEG vs nonDEG), P < 2.2e-16 (upDEG vs downDEG), P < 2.2e-16 (nonDEG vs downDEG). **d** Comparison of log2 fold changes in RPB1 occupancy between promoters (-1 to 1kb) and gene bodies (TSS+1kb to TES) in auxin-treated cells relative to control cells and distributions of the log2 fold changes at promoters and gene bodies. P value and Spearman correlation coefficient (rho) were calculated using the Spearman rank correlation test. **e** Representative genes showing altered Pol II occupancy at promoters and gene body. WINT4 gene (top) exhibited reduced RPB1 occupancy at the promoter but enhanced occupancy at the gene body. PCSK9 (middle) decreased RPB1 occupancy at both promoter and gene body, while ALS2CL (bottom) increased it at both regions.

This analysis revealed a positive correlation between them, indicating that RPB1 levels at promoters and bodies generally increased and decreased in the same direction (Figs. 1d and 1e). However, cohesin loss influenced RPB1 density along gene bodies far less strongly than at promoters. Moreover, at promoters, the distribution of changes shifted toward a decrease, while the changes at gene bodies were almost equally distributed between increase and decrease. These observations suggested that cohesin depletion mainly reduced Pol II binding at promoters, but the decrease was not directly transmitted to gene bodies, resulting in a minor effect on gene expression.

### Cohesin promotes Pol II pausing at highly paused genes

We observed that acute cohesin loss hardly affected gene expression but altered Pol II distribution along genes. At most genes, promoters exhibited more noticeable changes in Pol II occupancy, while milder changes were detected at gene bodies. These observations suggested that cohesin depletion affected not only Pol II binding at promoters but also Pol II pausing. To quantify the extent of the pausing and elongation, the pausing index (PI) was calculated as the ratio of Pol II density at promoters to that at gene bodies (Fig. 2a). We observed a significant reduction in PI for both upDEGs and nonDEGs upon cohesin depletion (Fig. 2b). However, the decrease in PI was not detected at downDEGs. The depletion of another component of cohesin, SMC1 successfully validated the reductions in PI (Figs. S2a-S2c).

**Fig. 2:**
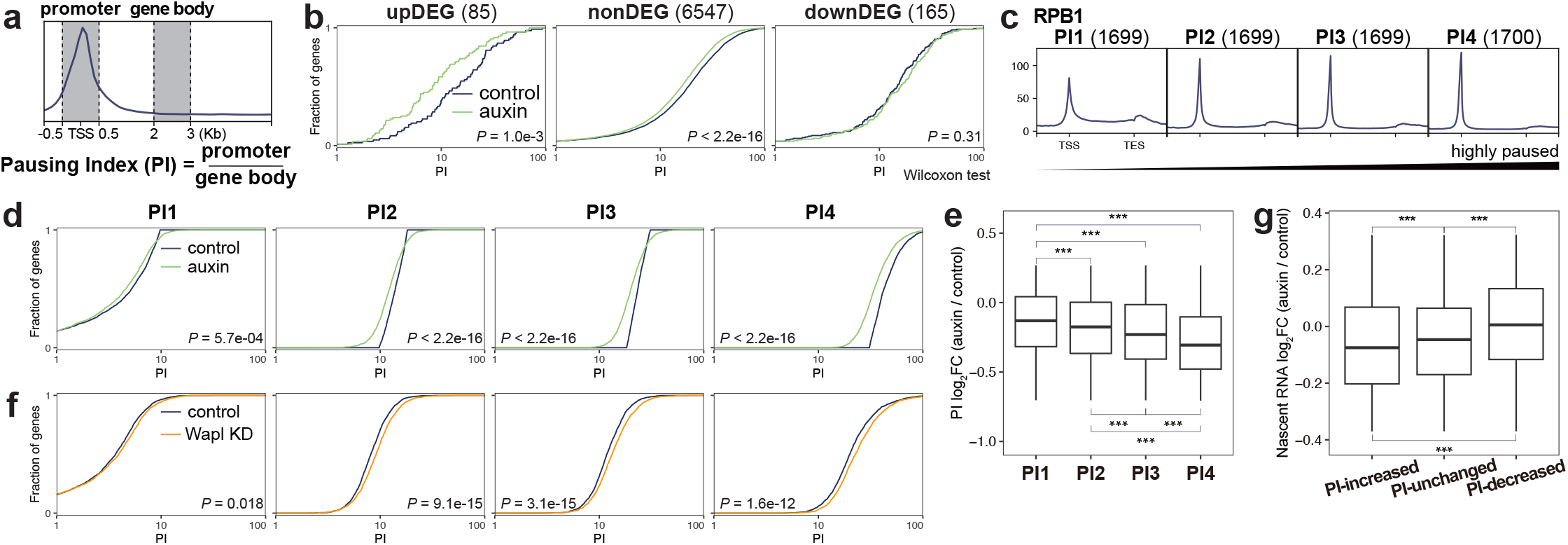
Cohesin promotes Pol II pausing at highly paused genes. **a** Pausing index (PI) is the ratio of Pol II occupancy in the promoters to that in the gene body. For genes longer than 2kb, PI was calculated as the ratio of read density in promoter (-500 to 500 bp) to that in gene body (2 to 3 kb). For genes less than 3kb, the gene body was defined as the region between 2kb from the TSS to TES. **b** Cumulative PI plot comparing control and auxin-treated cells for upDEGs, nonDEGs, and downDEGs. P values were calculated using the two-sided Wilcoxon rank-sum test. **c** Metagene plots of RPB1 from PI1 to PI4 genes and the number of the genes in parentheses. The analyzed genes were grouped by their PIs, and the groups were named PI1–PI4. PI1 represents genes with the lowest PIs and PI4 are genes with the highest PIs. **d** Cumulative PI plots comparing control and auxin-treated cells for PI1-PI4 genes. P values were calculated using the two-sided Wilcoxon rank-sum test. **e** Box plot showing the distribution of log2 fold change in PI upon auxin treatment at PI1-PI4 genes. A multiple Welch t-test was performed between every pair of gene groups. ***P[<[0.001. P = 8.9e-11 (PI1 vs PI2), P < 2.2e-16 (PI1 vs PI3), P < 2.2e-16 (PI1 vs PI4), P = 7.4e-3 (PI2 vs PI3), P < 2.2e-16 (PI2 vs PI4), P < 2.2e-16 (PI3 vs PI4). **F** Cumulative PI plots comparing control and Wapl KD cells for PI1-PI4 genes. P values were calculated using the two-sided Wilcoxon rank-sum test. **g** Box plot showing the distribution of log2 fold change in nascent RNA levels upon auxin treatment at PI-increased, PI-unchanged, and PI-decreased genes. A multiple Welch t-test was performed between every pair of gene groups. ***P[<[0.001. P = 7.6e-10 (PI-increased vs PI-unchanged), P < 2.2e-16 (PI-increased vs PI-decreased), P < 2.2e-16 (PI-unchanged vs PI-decreased).

To further investigate the effect of cohesin loss on different states of Pol II pausing, we divided genes into quartiles, from PI1 to PI4, based on their PI value (Fig. 2c). PI1 was the group with the lowest PI, whereas PI4 was the group with the highest PI value. Comparison between control and auxin-treated cells revealed a significant decrease in PI in all gene groups (Fig. 2d). However, PI4 showed the largest reduction in PI after cohesin loss, while PI1 exhibited the smallest decrease (Fig. 2e). SMC1-depleted cells reproduced the reductions in PI from PI1 to PI4, including the greatest decrease observed at PI4 (Figs. S2d and S2e). We also observed an overall negative correlation between PI itself and log2 fold-change of PI (Fig. S2f). These results indicated that cohesin depletion primarily reduced PI of highly paused genes. If cohesin contributes to stabilizing the paused state of Pol II, the increase in chromatin binding of cohesin would result in enhanced pausing. To test this, we knocked down cohesin-releasing factor Wapl, which dissociates cohesin from chromatin (Fig. S2g). We verified that the treatment increased Rad21 intensity at cohesin peaks (Fig. S2h). PI calculation revealed that Wapl knockdown significantly increased PI from PI2 to PI4, contrary to cohesin depletion, supporting our hypothesis (Figs. 2f and S2i).

Next, we classified genes based on changes in PI and found that cohesin loss increased PI at 1112 genes and decreased it at 3814 genes by more than 1.1-fold (Fig. S2f). The remaining 1872 were categorized as genes with unchanged PI. Consistent with our finding that a greater PI reduction was associated with highly paused genes, PI reduction was most frequently observed at PI4, while PI enhancement was most frequently seen at genes in PI1 (Fig. S2j). We observed that PI-increased genes reduced their nascent RNA levels upon cohesin depletion, while PI-decreased genes slightly enhanced the levels (Fig. 2g). This demonstrated that, as expected, an increase and decrease in PI were associated with the activation or repression of gene expression, respectively.

To study the impact of cohesin depletion on Pol II transition from pausing to elongation and its progression along gene bodies, we analyzed the binding profile of the elongating Pol II during synchronized transcription. We synchronized transcription at pausing step using flavopiridol, an inhibitor of Cdk9, and subsequently restarted productive elongation by washout of the inhibitor (Fig. S3a). ChIP-seq for RPB1 revealed the accumulation of Pol II at promoters in the presence of the inhibitor. At 10 and 15 min after the washout, we observed that the transcription wave shifted from promoters into gene bodies. Notably, even in the presence of flavopiridol, cohesin loss led to a reduction in RPB1 at promoters, consistent with our observations in the absence of the inhibitor (Fig. S3b). To observe the progression excluding the effect of the differential promoter binding of Pol II, we normalized RPB1 occupancy at each gene in cohesin-depleted cells by multiplying it with the ratio of RPB1 at the promoter in control to treated cells (Fig. S3c). This analysis revealed that PI-decreased genes longer than 40 kb exhibited promoted Pol II progression, while PI-unchanged genes also showed a mild enhancement in the progression. We measured the distance Pol II traveled using the island-based peak caller SICER and calculated the elongation rate based on the distance observed at 10 and 15 min (Fig. S3a and S3d).^34^ This measurement revealed that cohesin depletion enhanced elongation slightly at PI-increased genes and more substantially at PI-decreased genes (Fig. S3e). These results implied that cohesin loss affected Pol II dynamics, which we observed as an altered pausing degree during asynchronous transcription. When transcription was synchronized, the changed dynamics became evident as the enhanced Pol II progression. Taken together, our observations indicate that cohesin stabilizes Pol II pausing and restricts Pol II progression from promoters.

### Cohesin restricts Pol II release from pausing by inhibiting SEC recruitment to promoters

We observed a reduction in Pol II pausing upon cohesin depletion. Given the similarities between CdLS and CHOPS, it suggests that inactivation of cohesin and activation of SEC may have similar transcriptional effects. Since SEC is known to release Pol II from pausing, we investigated whether cohesin loss affected the genome-wide distribution of SEC, providing a potential explanation for the observed reduction in pausing. In addition to AFF4, a core subunit of SEC, we analyzed the genomic distribution of NELFCD, a subunit of NELF that stabilizes paused Pol II. ChIP-seq analysis revealed that cohesin depletion altered the binding of AFF4 and NELFCD along genes (Fig. 3a). To evaluate their abundance relative to Pol II, ChIP-seq signal of each factor was normalized to that of RPB1. This analysis revealed an increase in AFF4 and a decrease in NELFCD occupancy at promoters upon cohesin depletion (Fig. 3b). The enhanced AFF4 abundance at promoter still could be detected when Cdk9 was inhibited with 5,6-dichloro-1-β-D-ribofuranosyl-benzimidazole (DRB) (Fig. S4a). In contrast, treatment with flavopiridol, another inhibitor of Cdk9, rescued the reduction of NELFCD, indicating the dependency of NELF decrease on Cdk9 activity (Fig. S4b). Moreover, we also observed increases in AFF4 at gene bodies and Ser2P at promoters (Figs. 3c and S4c). Next, to test if SEC was involved in PI reduction observed after cohesin loss, we used KL-1, a SEC inhibitor. The presence of KL-1 suppressed the decrease in PI upon cohesin depletion (Fig. 3d and S4d). These results suggested that cohesin loss enhanced recruitment of SEC to Pol II at promoters before the activation of Cdk9, leading to an increase in SEC continuously associating with Pol II even during elongation (Fig. 3e). Furthermore, the increase in SEC would lead to enhanced phosphorylation of NELF and Pol II CTD mediated by Cdk9 within SEC, potentially promoting NELF dissociation from promoters and Pol II release from pausing simultaneously. As physical interactions between cohesin and SEC, along with Pol II and cohesin, were already reported, we speculate that cohesin inhibits SEC through these interactions.^29^

**Fig. 3:**
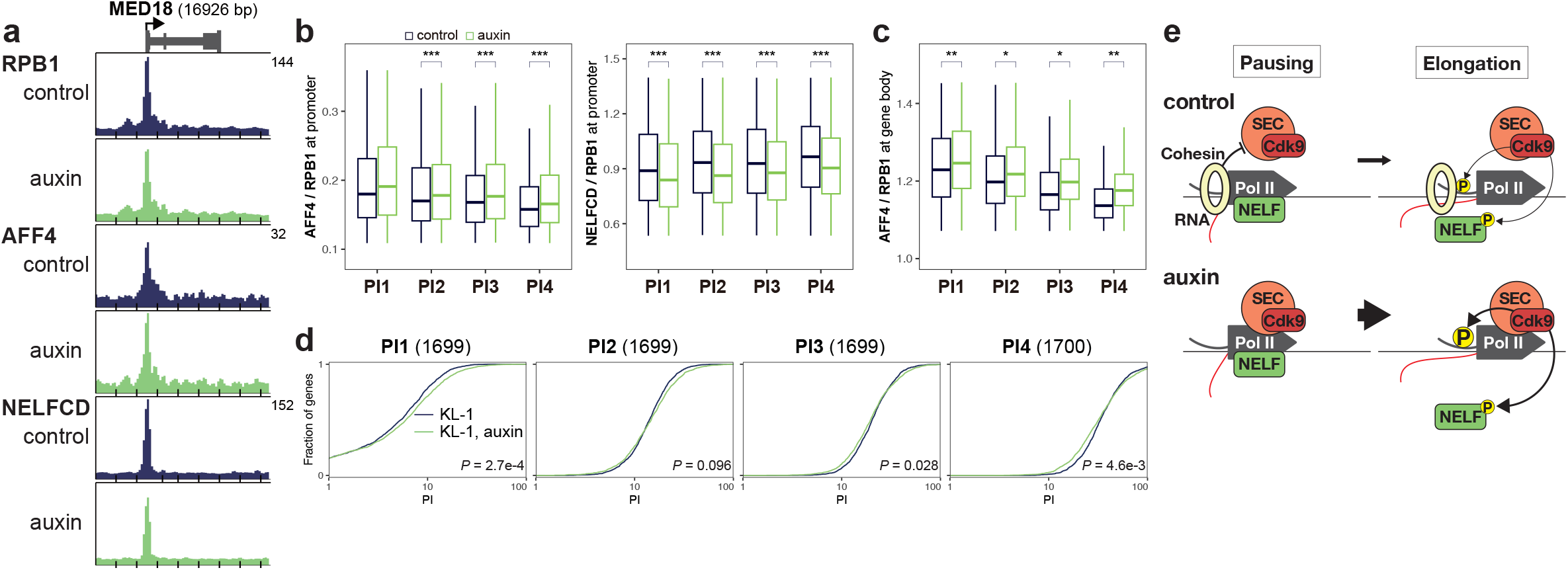
Cohesin restricts Pol II release from pausing by inhibiting SEC recruitment to promoters. **a** Representative gene showing increased AFF4 and decreased NELFCD occupancies at promoter. **b** Box plot showing the distribution of the relative amount of AFF4 and NELFCD to RPB1 at promoters (-500 to +500 bp) of PI-PI4 genes. P values were calculated using the two-sided Wilcoxon rank-sum test between control and auxin-treated cells. ***P[<[0.001. For AFF4, P = 5.1e-9 (PI2), P < 2.2e-16 (PI3), P < 2.2e-16 (PI4). For NELFCD, P =< 2.2e-16 (PI1), P =< 2.2e-16 (PI2), P < 2.2e-16 (PI3), P < 2.2e-16 (PI4). **c** Box plot showing the distribution of the relative amount of AFF4 to RPB1 at gene bodies (+500bp to TES) of PI1-PI4 genes. P values were calculated using the two-sided Wilcoxon rank-sum test between control and auxin-treated cells. *P[<[0.05, **P[<[0.01. P = 2.8e-3 (PI1), P = 2.4e-2 (PI2), P = 2.1e-2 (PI3), P = 3.1e-3 (PI4). **d** Cumulative PI plot comparing control and auxin-treated cells in the presence of KL-1 for PI1-PI4 genes. P values were calculated using the two-sided Wilcoxon rank-sum test. **e** Model for the role of cohesin in restricting SEC recruitment to promoters. In control cells, cohesin restricts SEC association with Pol II, inhibiting Cdk9-mediated phosphorylation of Pol II and NELF. In the absence of cohesin, SEC recruitment to Pol II at promoters is enhanced during the pausing step, leading to an increase in SEC continuously associating with Pol II during elongation. The increase in SEC leads to enhanced phosphorylation of NELF and Pol II CTD mediated by Cdk9 within SEC, promoting NELF dissociation from promoters and Pol II release from pausing simultaneously.

### Lowly paused genes exhibited intensive contacts with multiple H3K27ac-enriched enhancers

As cohesin is a major determinant of the 3D conformation structure of chromosomes, we investigated the effect of cohesin depletion on chromosome conformation by Micro-C. Our Micro-C contact maps generated from ∼1.99 billion pairwise interactions revealed finer and clearer chromatin structure than the Hi-C maps (∼2.60 billion reads) derived from a previous study that used the same cell line (Fig. 4a).^9^ Consistent with Hi-C, Micro-C revealed that rapid cohesin depletion changed the global distribution of chromosomal contact distances (Fig. S5a). TADs and chromatin loops were reduced and weakened by the depletion (Figs. S5b-S5e). In addition, Micro-C showed higher sensitivity for detecting chromatin loops than Hi-C, as already reported.^7,8,35^ Specifically, we identified 135539 Micro-C loops in control and 25989 in cohesin-depleted cells, which were approximately five times and 14 times the number of Hi-C loops, respectively (Figs. S5d, S5f, and S5g). Characterization of the loops that survived the depletion revealed that loops detected after cohesin loss were less likely to have cohesin peaks at their anchors and were relatively shorter compared to those in control cells (Figs. S5d and S5h). This can be interpreted that cohesin loss more effectively disrupted loops mediated by cohesin, which typically tend to be longer than other loops as previously reported (Fig. S5i and S5j).^36,13^

**Fig. 4:**
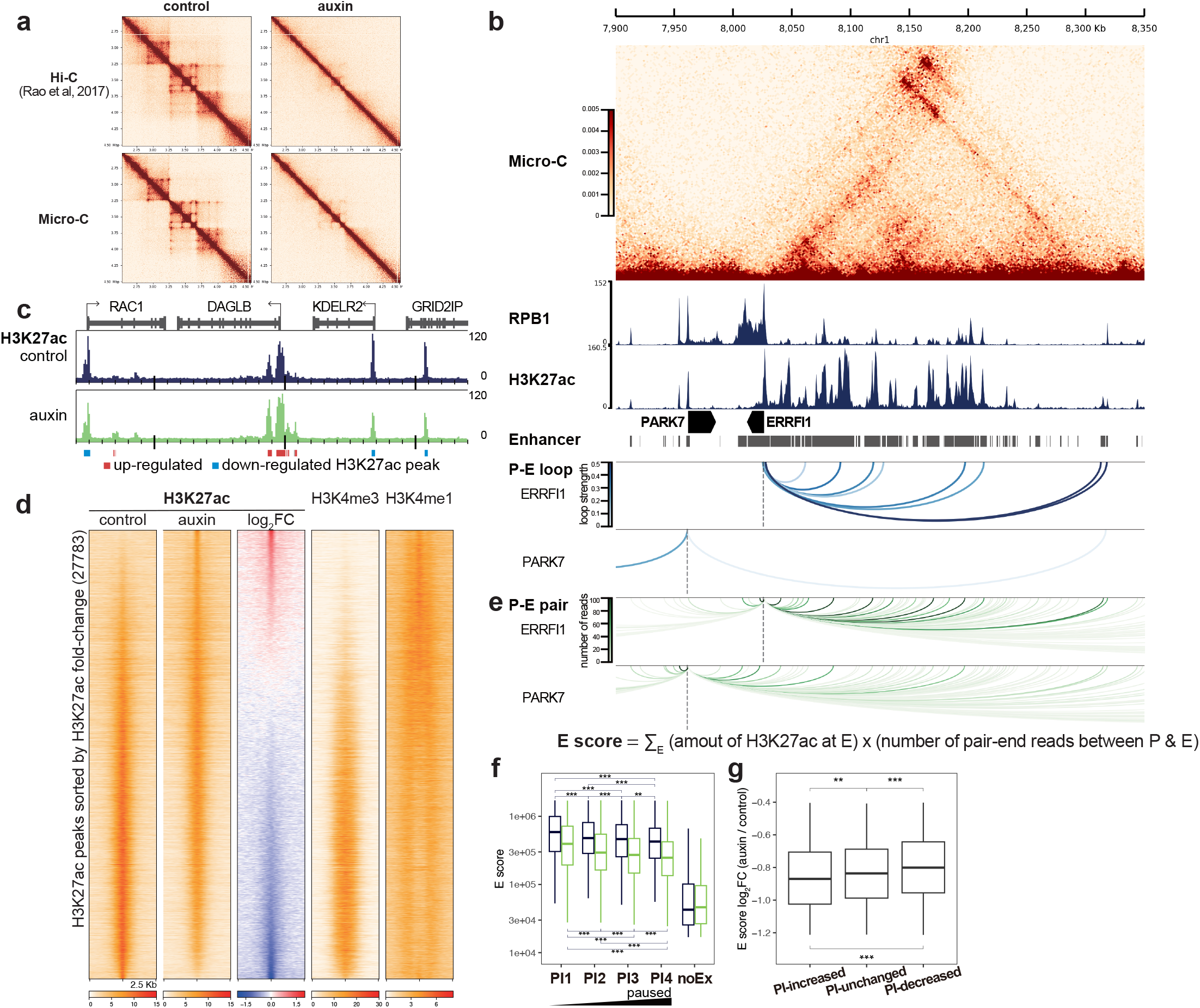
Lowly paused genes exhibited intensive contacts with H3K27ac-enriched enhancers. **a** Hi-C and Micro-C 5-kb resolution contact maps in the 2-Mbp region from chr18. Hi-C data were derived from Rao et al. (2017). **b** Micro-C 2-kb contact map aligned to ChIP-seq tracks of RPB1 and H3K27ac in the 4-Mbp region surrounding PARK7 (PI3) and ERRFI1 (PI1) genes. Gene models are shown in black with arrows indicating transcriptional orientation. Gray boxes indicate enhancer regions. Arcs represent Micro-C loops connecting promoters of the two genes and enhancers (P-E loops). The white-to-blue color scale reflects the loop strength. **c** Snapshots of spike-in normalized ChIP-seq signals of H3K27ac in chr7:6370000-6520000. Red and blue boxes indicate H3K27ac peaks up-regulated and down-regulated upon auxin treatment. **d** Heatmaps showing signals of H3K27ac, H3K4me3, and H3K4me1 at 27783 H3K27ac peaks sorted by log2 fold change in H3K27ac between control and auxin-treated cells. **e** E score for each gene was calculated by summing the number of pair-end reads connecting the gene and enhancer multiplied by the amount of H3K27ac at the enhancer. Arcs represent the pairs of promoters of PARK7 and ERRFI1 and enhancers (P-E pairs). The white-to-green color scale reflects the number of pair-end reads detected between the pair. **f** Box plot showing the distribution of E scores among PI1 to PI4 and noEx in control and auxin-treated cells. A multiple one-sided Wilcoxon rank-sum test was performed between every pair of gene groups. **P[<[0.01, ***P[<[0.001. For control cells, P < 2.2e-16 (PI1 vs PI2), P < 2.2e-16 (PI1 vs PI3), P < 2.2e-16 (PI1 vs PI4), P = 1.3e-8 (PI2 vs PI3), P < 2.2e-16 (PI2 vs PI4), P = 1.3e-3 (PI3 vs PI4). For auxin-treated cells, P < 2.2e-16 (PI1 vs PI2), P < 2.2e-16 (PI1 vs PI3), P < 2.2e-16 (PI1 vs PI4), P = 6.6e-7 (PI2 vs PI3), P = 2.8e-16 (PI2 vs PI4), P = 9.2e-4 (PI3 vs PI4). **g** Box plot showing the distribution of log2 fold change in E score at PI-increased, PI-unchanged, and PI-decreased genes. A multiple Welch t-test was performed between every pair of gene groups. **P[<[0.01, ***P[<[0.001. P = 1.2e-3 (PI-increased vs PI-unchanged), P < 2.2e-16 (PI-increased vs PI-decreased), P < 2.2e-16 (PI-unchanged vs PI-decreased).

Next, we aimed to characterize Micro-C loops connecting the promoters of PI1 to PI4 genes with enhancers (Fig. 4b). To identify these promoter-enhancer loops, enhancer regions were determined using ChIP-seq data of histone modifications. We segmented chromatin into eight chromatin states based on combinational histone modification marks using ChromHMM.^37,38^ Then, enhancers were defined as 67326 genomic regions with chromatin states marked by both H3K27ac and H3K4me1, indicative of active enhancer elements (see Method).39,40 The analysis revealed that over 20% of genes had no promoter-enhancer loops in control cells (Fig. S6a). This proportion increased to approximately 60% upon cohesin depletion. Genes in PI1 were connected to the most enhancers through the largest number of promoter-enhancer loops, and both the number of enhancers and loops gradually decreased from PI1 to PI4, both before and after cohesin loss (Figs. S6b and S6c). The intensity of loops between the most lowly paused PI1 genes and enhancers was slightly more intensive compared to other genes (Fig. S6d). Thus, analysis of promoter-enhancer loops suggested that lowly paused genes were more frequently and intensively connected with enhancers. However, not all genes were included in this analysis as we could not detect promoter-enhancer loops at every gene.

During analysis of histone modifications, we found up-regulated and down-regulated peaks of H3K27ac upon acute cohesin loss (Fig. 4c). Furthermore, the up-regulated and down-regulated peaks exhibited strong H3K4me1 and H3K4me3, respectively (Fig. 4d). H3K4me1 and H3K4me3 are well-established marks associated with enhancers and promoters individually.^39^ Pol II also increased and decreased at H3K27ac peaks upon cohesin loss in a similar way to H3K27ac, while H3K4me3 and H4K4me1 did not (Fig. S6e). This result was consistent with the observation that Pol II binding at the majority of promoters decreased after cohesin loss (Fig. 1b). Previous studies have shown that enhancer activities correlated with the intensity of H3K27ac at enhancers.^41,42,43,44^ Thus, this finding demonstrated the importance of accounting for H3K27ac redistribution when investigating the effect of cohesin depletion on promoter-enhancer communications.

To quantify regulatory signals from enhancers at individual gene level without relying on promoter-enhancer loops, we developed E score. E score is calculated as the sum of the paired-end read number multiplied by the strength of H3K27ac signal at the enhancers at each gene (Fig. 4e). This score utilized the H3K27ac signal as an indicator of enhancer strength and the number of the pair-end reads between promoters and enhancers as a parameter reflecting contact frequency. A previous study proposed a similar model and validated it with experimental data, supporting our idea that multiple enhancers additively regulate gene expression of their target genes.^45^ Every gene we selected for this study had at least one pair-end read connected to enhancers, which allowed us to calculate E score for all genes. We calculated E score for genes without any H3K4me3 peaks around TSS nor any transcripts (noEx) and found the lowest score compared to PI1 to PI4 genes, as expected (Figs. 4f and S6f). We observed that, in control cells, lowly paused PI1 genes had the highest E scores, while the most highly paused PI4 genes exhibited the lowest scores, apart from noEx. This result is consistent with our promoter-enhancer loop analysis, which showed that lowly paused genes were connected to more enhancers via greater in number and relatively intensive promoter-enhancer loops (Figs. S6b-S6d). Next, we compared PI and E score changes upon cohesin depletion. This revealed that PI-increased genes reduced E score more substantially, while PI-decreased genes exhibited a milder reduction (Fig. 4g). Thus, we observed the inverse relationship both between E score and PI itself, as well as between changes of the two parameters. These results suggested that the intensive interaction with robust and abundant enhancers would facilitate the efficient release of Pol II from pausing.

As the cohesin loss-mediated changes that would affect Pol II pausing and progression, we observed an increase in SEC binding along genes and a decrease in E score (Figs. 3b, 3c, and 4g). Each alternation is expected to promote Pol II release and progression and enhance the pausing, respectively. We propose that at PI-increased genes, the effect of reduced enhancer interactions overcame that of enhanced SEC activity, while the reverse was true at PI-decreased genes.

### Cohesin facilitates both Pol II binding at promoters and pausing simultaneously

We have shown that acute cohesin depletion impacted both promoter binding of Pol II and pausing (Figs. 1b and 2e). However, our gene expression analysis revealed only minor changes in the transcriptome upon the depletion. Here, we studied the relationship among Pol II promoter level, pausing, and gene expression. Comparison between changes in Pol II promoter binding and expression showed that about 90% of upDEGs exhibited increased or unchanged Pol II levels at promoters (Fig. S7a). In comparison, more than 90 % of downDEGs showed a reduction in the promoter level. Moreover, gene grouping according to PI changes revealed that about 80% of upDEGs were PI-decreased genes, while about 80% of downDEGs fell into PI-increased genes (Fig. S7b).

For a more detailed analysis, we classified genes based on a combination of Pol II promoter levels and PI. Nine new groups were generated by integrating the two gene groups of Pol II promoter levels and PI. Applying this grouping to all studied genes revealed that 35%, constituting the largest proportion, decreased both Pol II at promoters and PI (Fig. S7c). This aligns with our overall observation that cohesin loss inhibited Pol II promoter binding and pausing (Figs. 1b and 2e). Next, we applied this new categorization to upDEGs, nonDEGs, and downDEGs (Fig. 5a). Because 96% of genes were identified as nonDEGs, they displayed a similar distribution to that of all genes, with 36% exhibiting reduced promoter levels and PI. Compared to nonDEGs, upDEGs and downDEGs exhibited distinct proportions. Specifically, 39% of upDEGs displayed consistent promoter binding and decreased PI, while 44% of downDEGs reduced promoter binding and increased PI. Then, to further study how RPB1 promoter levels and PI impacted individual gene expression, we compared the changes of the three parameters at each gene (Fig. 5b). This analysis revealed the general trend that genes exhibiting increased promoter binding and reduced PI up-regulated their transcription, while genes with decreased Pol II at promoters and enhanced PI down-regulated transcription.

**Fig. 5:**
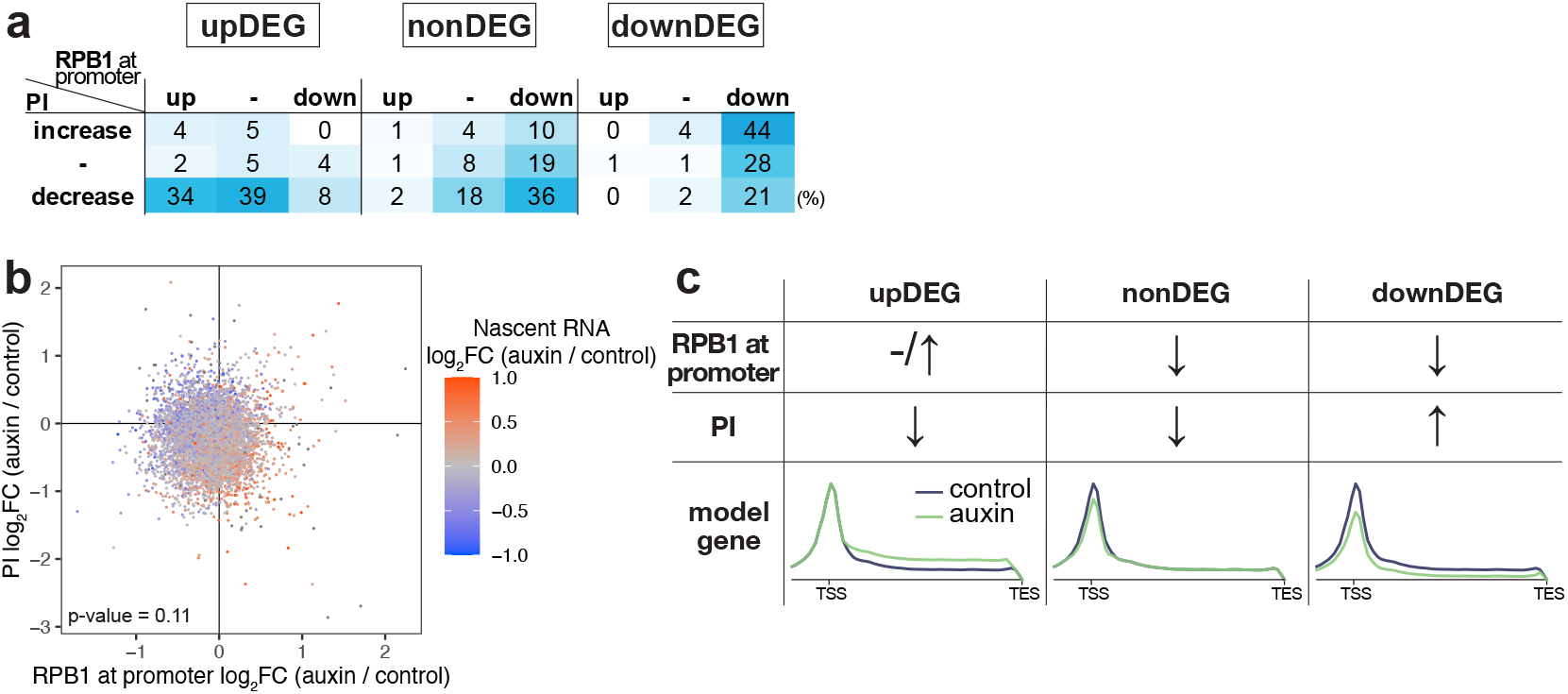
Cohesin facilitates both Pol II binding at promoters and pausing simultaneously. **a** Color scale table showing the proportions of the genes whose RPB1 promoter occupancy and PI were affected by auxin treatment. The percentages were calculated for upDEGs, notDEG, and downDEGs. **b** Comparison of log2 fold changes in RPB1 promoter occupancy (-1 to1kb) and PI in auxin-treated cells relative to control cells. The color of each dot corresponds to log2 fold change of EU signals of the gene. P value was calculated using the Spearman rank correlation test. **c** Model for the transactional effect of cohesin depletion. Auxin treatment reduced Pol II promoter binding and pausing. At most genes, the two contrary effects compensate for each other, leading to unchanged gene expression. At upDEGs, decreased pausing surpassed the reduction in Pol II at promoter, and vice versa at downDEGs.

Our observations suggested that cohesin loss primarily reduced Pol II loading at promoters and the pausing. The simultaneous reduction in both parameters is likely to result in unchanged gene expression due to a balance between them (Fig. 5c). Notably, we observed unaffected expression in the majority of genes upon cohesin loss. However, only a small subset of genes lost the balance and exhibited differential expression. Unchanged or increased promoter binding along with decreased pausing would lead to up-regulated transcripts, while enhanced pausing with reduced promoter levels is expected to result in down-regulated expression.

## Discussion

We have shown that acute cohesin loss reduces Pol II pausing at the majority of genes. As a candidate mechanism contributing to pausing reduction, we observed an increase in SEC relative to Pol II at promoters and gene bodies after cohesin loss. Notably, in both CdLS and CHOPs syndrome cell lines, we also observed enhanced SEC abundance at expressed genes (unpublished data). Our previous work demonstrated similar dysregulated gene expressions in CdLS and CHOPS, which is considered the direct cause of the deseases.^29^ These observations suggest that this enhanced SEC occupancy along genes is a shared mechanism between CdLS and CHOPS, contributing to the dysregulated transcriptome. We previously demonstrated the physical interaction between cohesin and SEC, as well as between cohesin and Pol II.^29^ One possible mechanism for limiting SEC recruitment by cohesin is that cohesin acts as a physical barrier, inhibiting their interaction between SEC and Pol II in control cells. Further investigation is needed to uncover the mechanism by which cohesin loss enhances SEC recruitment.

The majority of genes showed decreased pausing after cohesin depletion. On the other hand, small portion of genes exhibited enhanced pausing even though SEC binding increased, suggesting another cohesin-related regulatory mechanism of pausing. Then, we investigated promoter-enhancer interactions and found that these genes showed a more substantial reduction in positive signals from enhancers (E score). We observed negative correlation between pausing and E score, which means lowly paused genes exhibit higher E scores. Moreover, previous work shows that depletion of enhancer regions enhance pausing of their target genes.^46^ Taken together, these results imply that enhancers can mediate efficient Pol II release from the paused state. Based in these observations, we propose that cohesin regulates Pol II pause and release both by limiting SEC recruitment to promoters and by mediating promoter-enhancer interactions. In this model, an increase in SEC binding observed after cohesin loss genes will result in reduced pausing, while a decrease in E score will lead to enhanced pausing. We speculate that small part of genes showed the enhanced pausing because the effect of reduced E score outweighed that of enhanced SEC activity. Increased H3K27ac after hormone stimulation or histone deacetylase inhibition has been reported to enhance Pol II release from the paused state.^47,48^ These findings partially support our model, though they do not specifically address promoter-enhancer interactions. To date, very few studies have investigated which transcriptional step promoter-enhancer interactions modulate. Thus, our findings can contribute to a better understanding of the involvement of promoter-enhancer contacts in transcriptional regulation.

H3K27ac and Pol II exhibited similar decreases at promoters and increases at enhancers upon cohesin loss. The genomic colocalization with cohesin is not associated with the increase or decrease of the two factors. We speculate that the interaction between promoters and enhancers is involved in the redistribution. A model where Pol II is recruited to enhancers and then transferred to promoters of the target genes has been proposed.^49,50^ Moreover, artificial tethering of histone acetyltransferase (HAT) domain of p300 to enhancers enhanced H3K27ac not only at the enhancers but also at the target genes.^51^ Based on these findings and our observations, we propose that enhancers can supply Pol II and H3K27ac to the target promoters through promoter-enhancer interactions. Loss of promoter-enhancer interactions upon cohesin depletion possibly inhibits the transfer from enhancers to promoters, allowing these factors to accumulate at enhancers and decrease at promoters.

We observed simultaneous decreases in Pol II promoter binding and Pol II pausing at many genes upon cohesin depletion. If the observed reduction in pausing was caused by decreased Pol II promoter binding, genes with a substantial decrease in the promoter levels should have exhibited the most prominent reduction in pausing. However, we found no significant correlation between changes of Pol II promoter binding and pausing. This result suggests that cohesin loss independently affected Pol II promoter binding and pausing status. Moreover, synchronization of transcription revealed that cohesin loss altered Pol II, which also validated the effect of cohesin on transcription elongation.

The connection between pausing and release and co-transcriptional events like splicing, 5’-capping, and termination has been proposed.^33,52,53,54,55,56^ Combinations of mutations in cohesin subunits and splicing factors, such as *SRSF2* and *SF3B1I*, have been reported to cause acute myeloid leukemia.^57,58^ These raised the possibility that cohesin may regulate these events by promoting RNAPII pausing. Thus, future investigation will be to assess the effect of the loss on transcribed RNA itself.

## Method

### Cell culture

HCT-116-RAD21-mAID-mClover has been reported previously.^59^ HCT116 cells were cultured in McCoy’s 5A medium containing 10% FBS, Penicillin-Streptomycin-l-Glutamine Solution (Wako) at 37°C with 5% CO2. Treatment with 0.5 mM indole-3-acetic acid (IAA, SIGMA) for 3 hr was performed to deplete Rad21. For inhibition of SEC, cells were treated with 20 μM KL-1 (MedChemExpress) for the last 2 hr of the 3-hr auxin treatment. For inhibition of Cdk9, cells were treated with 100 μM 5,6-Dichlorobenzimidazole 1-β-D-Ribofuranoside for 3 hr.

### siRNA knockdown of Wapl

RNAi was performed using Lipofectamine RNAiMax (Invitrogen) with a final RNA duplex concentration of 50 nM according to the manufacturer’s instructions. Two days after transfection, cells were collected. The sequence of siRNA was 5’-UCUCCUGCUCGUUAGAAGUAAGGGb-3’.

### Generation of HCT-116 SMC1A-mAID

HCT116 CMV-OsTIR1 SMC1A-mAID was generated as previously reported.^59,60^ HCT116 CMV-OsTIR1 cells were co-transfected with SMC1A CRISPR and SMC1A-mAID-Hygro donor vectors. After selection with 100 μg/mL HygroGold (Invivogen), single colonies were isolated and recombination-mediated knock-in of mAID at the SMC1A allele was confirmed by genomic PCR and western blotting.

### Cell cycle profile analysis

The cells were suspended in PBS and fixed with 70% ethanol. The fixed cells were washed with PBS twice, treated with RNaseA (50 μg/mL), and stained with 50 μg/ml propidium iodide. DNA content per cell was measured by flow cytometer (BD Accuri C6 plus).

### ChIP-seq

ChIP was performed as described in Sakata et al.^61^ HCT116 and C2C12 cells were crosslinked with 1% formaldehyde for 10 min, quenched with 125 mM glycine for 5 min, and washed with PBS. After collection by centrifugation, the fixed cells were frozen and stored at -80°C. The thawed cells resuspended with LB1 Buffer (50 mM HEPES-KOH (pH 7.5), 140 mM NaCl, 1 mM EDTA, 10% Glycerol, 0.5% NP-40, 0.25% Triton-X, 1 mM PMSF, 10 mM DTT, 1x cOmplete EDTA-free (Roche)) and incubated on ice for 10 min. After collection by centrifugation, the pellet was resuspended with LB2 Buffer (20 mM Tris-HCl (pH 7.5), 200 mM NaCl, 1 mM EDTA, 1 mM PMSF, 1x cOmplete EDTA-free) and incubated on ice for 10 min. After centrifugation, the pellet was resuspended with LB3 Buffer (20 mM Tris-HCl (pH 7.5), 150 mM NaCl, 1 mM EDTA, 1% Triton-X, 0.1% Na-Deoxycholate, 0.1% SDS, 1x cOmplete EDTA-free) and incubated on ice for 10 min. Then, we repeated the centrifugation and resuspension with LB3 Buffer. For shearing chromatin, sonication was performed for 12 sec six times with amplitude setting at 17% of the maximum amplitude using Sonifier 250D. Between each cycle, chromatin was collected by centrifugation. After the step of sonication, cell debris was removed by centrifugation at 19000 g for 15 min. 50 mL of Dynabeads Protein A or Dynabeads Protein G (Invitrogen) was washed with 5 mg/mL BSA/PBS twice and resuspended with BSA/PBS. 4 μg antibody was added to the beads, and the mixture was rotated for 3 hr at 4°C. The beads were washed with BSA/PBS twice and with LB3 Buffer once and resuspended with LB3 Buffer. Chromatin from HCT116 and from C2C12 cells were mixed and incubated with antibody beads for 14[hr at 4°C. The beads was washed with LB3 Buffer once, with LB3 High Buffer (20 mM Tris-HCl (pH 7.5), 500 mM NaCl, 1 mM EDTA, 1% Triton-X, 0.1% Na-Deoxycholate, 0.1% SDS) twice, with RIPA wash Buffer (50 mM HEPES-KOH (pH 7.4), 0.25 M LiCl, 1 mM EDTA, 0.5% Na-Deoxycholate, 1% NP-40) three times and with TE50 Buffer (50 mM Tris-HCl (pH 7.5), 10 mM EDTA) once. Chromatin bound to beads was eluted with Elution Buffer (50 mM Tris-HCl (pH 7.5), 10 mM EDTA, 1% SDS). The eluates were incubated at 65°C for 6 hr to reverse crosslinks and then treated with RNaseA and then with proteinase K. The samples were purified with QIAquick PCR purification kit (QIAGEN). The library for sequencing was prepared with NEBNext DNA Library Prep Master Mix Set for Illumina and NEBNext Multiplex Oligos for Illumina (New England Biolabs). The library was sequenced on Hiseq 2500 (Illumina) or Nextseq 2000 (Illumina).

The following antibodies were used: Rpb1 NTD (D8L4Y) antibody (Cell Signaling Technology, #14958), RAD21 (D213) Antibody (Cell Signaling Technology, #4321), Purified Rabbit Anti-GFP, Anti-GFP antibody (Torrey Pines Biolabs, TP401), Anti-Phospho RNA Polymerase II CTD (Ser2) (MBL International, MABI0602), anti-acetyl Histone H3 (Lys27) antibody (MBL International, MABI0309), anti-trimethylHistone H3 (Lys4) antibody (MBL International, MABI0304), anti-monomethyl Histone H3 (Lys4) antibody (MBL International, MABI0302), Rabbit anti-MCEF Antibody (Bethyl, A302-539A), TH1L (D5G6W) Rabbit mAb (Cell Signaling Technology, #12265).

### ChIP-seq alignment and data analysis

Sequenced reads were aligned to hg38 and mm10 using bowtie^62^ (version 1.2.3) with “-n2-m1” parameters. The removal of redundant reads and the normalization of the read number for each sample were performed with parse2wig^63^ (version 3.7.2). For peak-calling, we used drompa_peakcall^63^ (version 3.7.2) in “PC_SHARP” mode, and for visualization, we used drompa+ PC_SHARP^64^ (version 1.17.2) and deeptools^65^ (version 3.5.1.)

### Quantitative PCR

qPCR of ChIP samples was performed using real-time PCR systems 7500 and StepOnePlus (Life Technologies) and KAPA SYBR Fast qPCR kit (KAPA Biosystems). The primer sequences were as follows:

cohesin site #1 forward primer: 5’-GGCCTAGAAAACACGTCTACAG-3’

cohesin site #1 reverse primer: 5’-TCCACCAAGGGACAAACAGAG-3’

cohesin site #2 forward primer: 5’-TGCGGGTACACTTTGAGAACC-3’

cohesin site #2 reverse primer: 5’-GTGCCACCTAGTGGTATTTCCT-3’

cohesin site #3 forward primer: 5’-GGCCTAGAAAACACGTCTACAG-3’

cohesin site #3 reverse primer: 5’-ATCCTGACACTGCTGGGAATG-3’

Negative Control forward primer: 5’-ATGAGTCTGGGATTTGGCTGG-3’

Negative Control reverse primer: 5’-TCTCAGGGGAAGGAGAGAGTC-3’

### EU-seq

For labeling of nascent transcripts, HCT116 cells and S2 cells were incubated in 0.5 mM 5-ethynyl uridine (Thermo Fisher) for 20 min and 0.2 mM for 2 hr. Total RNA from HCT116 cells and drosophila S2 cells was isolated with Trizol (Invitrogen) and Nucleospin RNA II (Macherey-Nagel) following the manufacturer’s instructions. rRNA was depleted from total RNA from HCT116 cells with NEBNext rRNA Depletion Kit (NEB). mRNA was purified from total RNA from S2 cells with Dynabeads mRNA Purification Kit (Invitrogen). 200 ng of rRNA-depleted RNA of HCT116 cells and 20 ng of mRNA of S2 cells were mixed, and EU-labelled RNA was isolated with Click-iT Nascent RNA Capture Kit (Thermo Fisher). Single-strand cDNA was synthesized from the isolated RNA by SuperScript VILO cDNA synthesis kit (Thermo Fisher). Second-strand cDNA synthesis and the subsequent steps to create library were conducted with NEBNext Ultra II Directional RNA Library Prep Kit for Illumina and NEBNext Multiplex Oligos for Illumina (New England BioLabs). The library was sequenced on Nextseq 2000 (Illumina).

### EU-seq alignment and data analysis

Low-quality reads were removed with Trimmomatic^66^ (version 0.39) using the parameters LEADING:30 TRAILING:30 SLIDINGWINDOW:4:15 MINLEN:20.

Single-end 65-bp reads were aligned to dm6 with Bowtie2^67^ (version 2.4.5) in “--very-sensitive-“ mode and the unmapped reads were aligned to hg38 in the same mode. The removal of redundant reads and the normalization of the read number for each sample were performed with parse2wig (version 3.7.2). For visualization, we used deeptools.

### Micro-C

HCT116 cells were dissociated into single cells with TrypLE Express (Thermo Fischer Scientific) and washed with PBS once. The washed cells were crosslinked with 1% formaldehyde for 10 min and quenched with 125 mM glycine for 5 min. After washing with PBS, the cells were crosslinked with 3 mM DSG (Thermo Fisher) for 40 min and quenched with 400 mM glycine for 5 min. The crosslinked cells were washed with PBS and frozen in aliquots of 5 x 10^6^ cells. Frozen cells were resuspended in Micro-C Buffer #1 (50 mM NaCl, 10 mM Tris-HCl pH 7.5, 5 mM MgCl2, 1 mM CaCl2, 0.2% NP-40, 1x cOmplete EDTA-free (Roche)) and incubated on ice for 20 min. Cells were collected by centrifugation and resuspended with Micro-C Buffer #1 again. The treatment with MNase (NEB) at 37°C for 10 min fragmented chromatin. The reaction was stopped by EDTA and incubation at 65°C for 10 min. The chromatin was collected by centrifugation and washed with Micro-C Buffer #2 (50 mM NaCl, 10 mM Tris-HCl pH 7.5, 10 mM MgCl2). To generate 5’ overhangs, the chromatin was resuspended with 50 mL end-repair mix (5 μL of 10 x NEBuffer 2.1, 2 μL of 100 mM ATP, 2.5 μL of 100 mM DTT, 8 μL of Large Klenow Fragment (NEB), 2 μL of T4 Polynucleotide Kinase (NEB).) and incubated at 37°C for 15 min. The DNA overhangs were filled with biotinylated nucleotides by addition of 100 μL Fill-in master mix (10 μL of 1 mM biotin-14-dATP (JBS), 10 μL of 1 mM biotin-11-dCTP (JBS), 1 μL of 10 mM dGTP (Thermo Fisher), 1 μL of 10 mM dTTP (Thermo Fisher), 10 μL of 10x NEBuffer 2.1). After incubation at 65°C for 20 min, the reaction was stopped by EDTA. The chromatin was collected with centrifugation and washed with 1x Ligase Reaction Buffer (NEB). For proximity ligation, the chromatin was resuspended with 500 μL Micro-C ligase mix (50 μL of 10x NEB Ligase buffer, 2.5 μL of 200x BSA (NEB), 5 μL of NEB T4 Ligase (NEB)) and incubated at RT form 2.5-3 hr. For removal of biotin-dNTP from un-ligated ends, collected chromatin by centrifugation was resuspended with 100 μL of exonuclease mix (1x NEBuffer 1 (NEB) and 2 μL of 100 U/mL NEB Exonuclease III (NEB)) and incubate at 37°C for 15 min. For deproteination and reverse crosslinking, 15 μL of Proteinase K (50 mg/mL) and 13 μL of 10% SDS (Invitrogen) were added and incubated at 55°C for 2h and at 65°C overnight. Ligated DNA was purified with QIAquick PCR purification kit (QIAGEN) and eluted in 110 μL of EB. Size selection of DNA was performed by AMPure XP beads (Beckman Coulter). 40 μL of beads were added to each reaction and incubated for 5 min. Using a magnetic stand, the clear solution was transferred to a fresh tube, and 20 μL of beads were added. After incubation for 5 min, the supernatant was removed, and beads were washed with 70% ethanol without mixing. Beads were left to dry for 5 min and DNA was eluted with 50 μL of EB. For biotin pull-down, we used Dynabeads™ MyOne™ Streptavidin C1 beads. 5 μL beads were washed twice with 300 mL 1x Tween Washing Buffer (1x TWB; 5 mM Tris-HCl (pH 7.5), 0.5 mM EDTA, 1 M NaCl, 0.05% Tween 20) and suspended in 150 μL 2x Binding Buffer (2x BB; 10 mM Tris-HCl (pH 7.5), 1 mM EDTA, 2 M NaCl). 100 μL H2O and 150 μL washed beads in 2x Binding Buffer were added to the sample and incubated rotation at RT for 20 min. Beads were washed with 1x TWB twice and with EB once. End repair and adaptor ligation was performed on beads using NEBNext Ultra II DNA Library Prep Kit for Illumina (NEB) according to manufacturer’s protocol. Then, beads were washed with 1x TWB twice and with EB once and resuspended with 15 μL of EB. DNA was amplified by PCR and DNA purification was performed using NEBNext Ultra II DNA Library Prep Kit, NEBNext® Multiplex Oligos for Illumina and AMPure XP beads following the manufacturer’s instructions. The library was sequenced on NovaSeq 6000 (Illumina).

### Micro-C alignment and data analysis

The sequenced reads were aligned to hg38 with bwa mem^68^ in “-5SP -T0 -t16” parameters. Identification of ligation junctions, sorting of reads, removal of PCR duplication and generation of pairs files were performed with pairtools^69^ (version 0.3.0). From the pairs files, matrix files were made by Juicer^70^ (version 1.22.01) at 5k, 2k and 1k resolution.

Chromatin loops were identified using Juicer Tools HiCCUPS and Mustache^71^ (version 1.2.0) at 5k, 2k and 1k resolution. Loops called by HiCCUPS and Mustache at various resolutions were combined to generate a set of Micro-C loops. TADs were identified using Juicer Tools Arrowhead at 25k resolution. The intensity of Micro-C loops were quantified using chromosight^72^ (version 1.6.1).

For visualization, we used HiCExplorer^73^ to obtain contact maps and coolpup.py^74^ to generate aggregate plots.

### Selection of genes for analysis

We selected 6797 genes with H3K4me3 peaks within 5kb from their TSSs, with RPKM of RPB1 ChIP-seq > 1, > 2kb in length, and >1 kb away from any other genes.

### Identification of up- or down-regulated H3K27ac peaks

We identified H3K27ac peaks in control and auxin-treated cells, using drompa_peakcall in “PC_SHARP -binsize 100 -pthre_enrich 0.001 -pthre_internal 0.001 -norm 0” parameters. H3K27ac peaks detected in control and auxin-treated cells were merged to generate the total set of H3K27ac peaks. H3K27ac signal in control and auxin-treated cells were compared by binominal test using drompa_draw CI at the peak sites. Using p.adjust in R, FDR was calculated from the p-value for each peak. We identified increased peaks with FDR < 0.05 as up-regulated peaks and decreased peaks with FDR < 0.05 as down-regulated peaks.

Identification of genes with up- or down-regulated RPB1 signal at promoters The genes analyzed in this study should have H3K4me3 peaks within 5kb of their TSSs, an RPKM value of RPB1 greater than 1, a gene length longer than 2kb, and be at least 1 kb away from other genes. RPB1 signal in control and auxin-treated cells were compared by binominal test using drompa_draw CI at regions within 1 kb from TSSs. Using p.adjust in R, FDR was calculated from the p-value for each promoter. We identified increased promoters with FDR < 0.05 as up-regulated promoters and decreased promoters with FDR < 0.05 as down-regulated promoters.

### Calculation of pausing index (PI) calculation

For genes analyzed in this study, RPB1 occupancy on the promoter (within 0.5 kb from TSS) and on gene body (+2 kb to +3 kb) was determined using multiBigwigSummary from deeptools. Pol II density on the promoter and gene body was calculated as the averaged RPB1 signals from two replicates. PI was calculated by dividing the Pol II density on the promoter by the Pol II density on the gene body. Genes with more than a 1.1-fold increase or decrease in PI were classified as PI-increased and PI-decreased genes, respectively.

### Identification of differentially expressed genes

We generated the read count table for the genes listed in the Ensembl annotation (Homo_sapiens.GRCh38.103.gtf). The number of EU-seq reads mapped to each gene was counted using drompa_draw CI. Differentially expressed genes were identified by the R Bioconductor package edgeR^75^ (version 4.2.2) using LRT (likelihood ratio test) methods (FDR < 0.05).

### Synchronization of transcription with Cdk9 inhibitor

For the last 1 hr of the 3-hr auxin treatment, cells were treated with 1 mM flavopiridol (Santa Cruz). After the treatment with flavopiridol, cells were washed with PBS twice and culture media once. Following an incubation with fresh media for 10 or 15 min, the cells were fixed for ChIP-seq. Cells for the 0 min time point were fixed after 1-h treatment with flavopiridol.

### Analysis of Pol II progression

We selected genes that were longer than 40 kb, had only one RPB1 peak at their promoter regions (TSS+/- 1kb) in the presence of flavopiridol, and showed an increased RPB1 signal in their gene bodies after the washout of flavopiridol. RPB1 signals of auxin-treated cells at 10 and 15 min after washout were normalized by multiplying it with the ratio of RPB1 at the promoter (TSS+/- 1kb) in control to auxin-treated cells. The ratio was calculated at each gene using multiBigwigSummary from deeptools, and the normalized profile was generated using computeMatrix from deeptools with the“- scale” option.

### Calculation of elongation rate

The distances Pol II traveled were determined by peak calling with SIDER in “-s hg38 - w 500 -g 6000” parameter.^34^ We selected genes longer than 40 kb that had SIDER- called peaks around their promoters (TSS +/- 1 kb) and no peaks at their gene bodies (TSS +10k-40k) at 0 min. In addition, at 10 and 15 min, the genes should exhibit a single peak originating from their promoters and elongating in a time-dependent manner. We defined the distance as the length from TSS to the end of the called peak. The elongation rate was calculated as the distance divided by the time after washout.

### Calculation of E score

We defined enhancer regions using ChromHMM 8-state model.^38^ As input, we used our ChIP-seq data of H3K27ac, H3K4me1, H3K4me3, H3K36me3 and Rad21, in addition to public data for CTCF, H3K27me3, H3K79me2, H3K9ac, H3K9me2, H3K9me3, H4K20me1, DNase and ChIP input. 67326 segments included in the cluster exhibiting strong H3K4me1 and H3K27ac were identified as enhancers in this study. The size of these enhancer regions was expanded by 1 kb in each direction. H3K27ac occupancy at these enhancer regions was calculated using the multiBigwigSummary from deeptools. For each gene, we extracted all contacts anchoring its promoter (TSS +/- 1 kb) at one end and enhancers at the other end using bedtools pairtobed. The E score of the gene was calculated as the sum of the number of contacts detected with an enhancer multiplied by the H3K27ac occupancy at the enhancer.

### Statistical analysis

Graphs were generated using R and Microsoft Office Excel 2023. Statistical significance was determined by a multiple Walch t-test, a two-sided Wilcoxon rank-sum test, a one-sided Wilcoxon rank-sum test, and the Spearman rank correlation test. *P*[<[0.05 was the criterion used to represent a significant difference.

### Data availability

FASTQ files and processed data were deposited in GEO with the accession ID GSE260852, GSE260853, and GSE260854.

### Code availability

All code is available upon request.

## Acknowledgment

We would like to thank all members of Shirahige Laboratory for discussions. This work was supported by JST-CREST Grant Number JPMJCR18S5, grant-in-Aid for Transformative Research Areas (A) Grant Number 20H05940 and 20H05933, AMED-ASPIRE-A, JSPS Grant-in-Aid for Scientific Research (S) Grant Number 20H05686, and AMED BINDS grant number 22ama121020j0001.

## Author information

### Contributions

S.T., T.S., M.B., and K.S. conceived the concept. T.M. and M.T.K generated the cell lines. S.T., A.Y., and M.B. performed the experiments. S.T. analyzed data and wrote the manuscript. S.T., T.S., M.B., and K.S. reviewed and edited the manuscript.

### Ethics declaration

Competing interests

The authors declare no competing interests.

