## Supplemental figures for "Cohesin regulates promoter-proximal pausing of RNA Polymerase II by limiting recruitment of super elongation complex"

#### Fig. S1

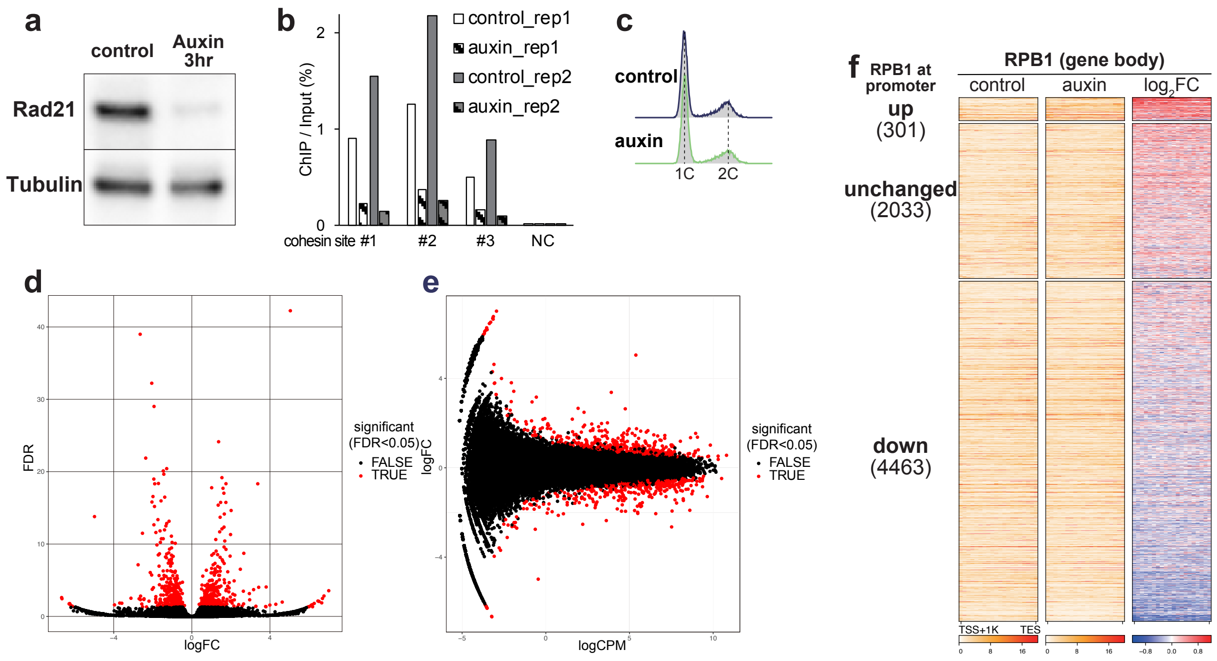


**Fig. S1: Cohesin regulates the expression of a subset of genes while promoting Pol II binding at the majority of promoters (related to Fig. 1)**

**a** Confirmation of cohesin depletion by immunoblotting with polyclonal antibody against Rad21. The band intensity of Rad21 normalized to Tubulin was reduced to 22% after auxin treatment for 3 hr.

**b** Confirmation of cohesin depletion by ChIP-qPCR at three Rad21 peaks and a negative control (NC) site. An antibody against GFP was used to detect Rad21 fused with mini auxin-inducible degron and mClover at the C-terminus.

**c** Histograms of DNA content. Auxin treatment for 3 hr did not affect the cell cycle progression.

**d** Volcano plot showing changes in nascent transcription (EU-seq) after auxin treatment. n = 2 biological replicates. FC, fold change. Differentially expressed genes (DEGs) were identified using the edgeR package. p-values were calculated using the likelihood ratio test. FDR was calculated by adjusting the p value for multiple testing using the Benjamini–Hochberg method. Among the genes, we used those with H3K4me3 peaks within 5kb of their TSSs, with RPKM $\geq 1$, > 2kb in length and >1kb away from the other genes.

**e** MA plot showing changes in nascent transcription (EU-seq) after auxin treatment. n = 2 biological replicates. CPM, counts per million mapped reads.

**f** Heatmaps showing signals of RPB1 in control and auxin-treated cells at gene bodies (TSS+1kb to TES) of genes with up-regulated, unchanged, and down-regulated RPB1 promoter occupancy upon auxin treatment. Gene grouping and sorting orders are identical to those in Fig. 1b.

#### Fig. S2

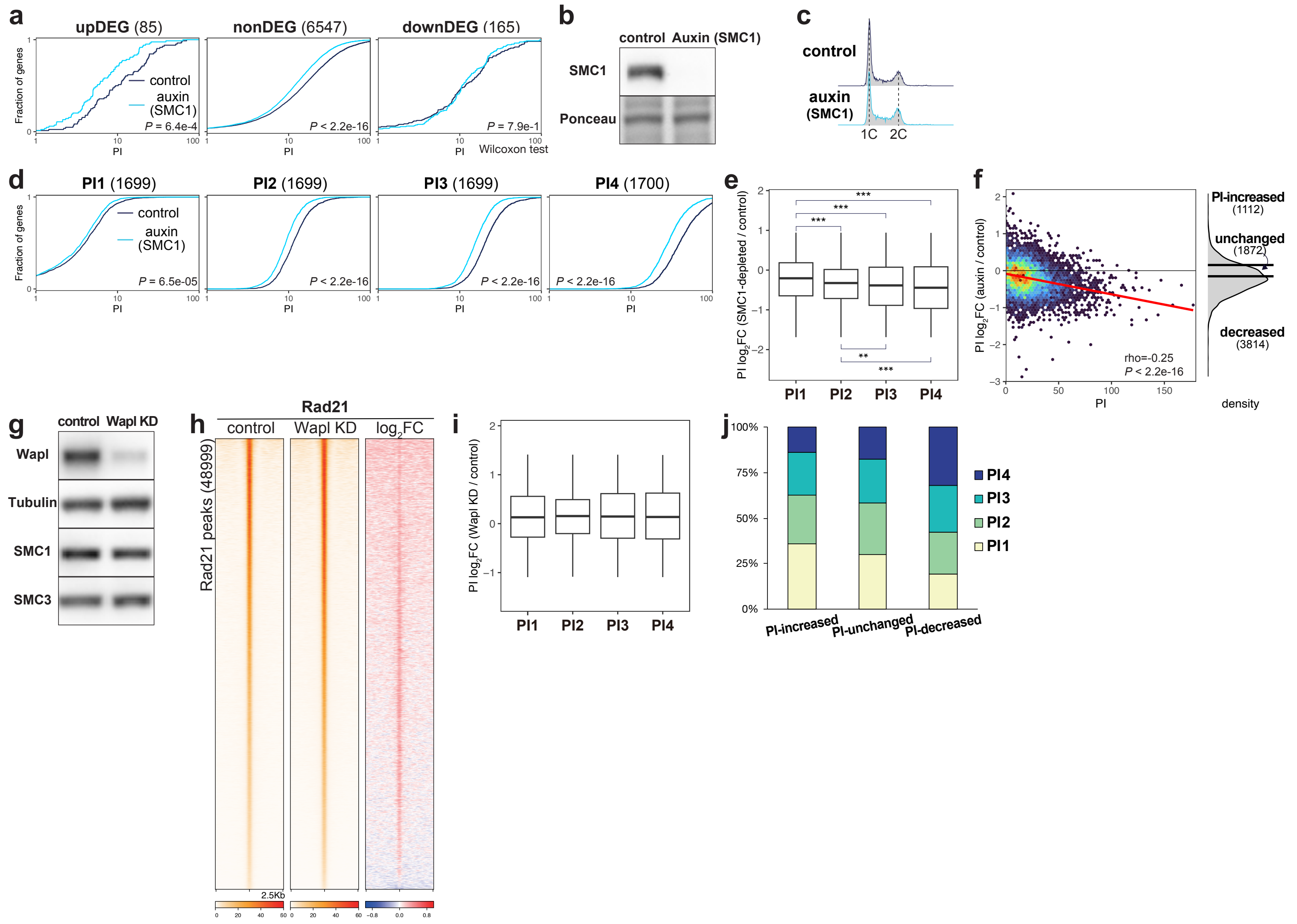


**Fig. S2: Cohesin promotes Pol II pausing at highly paused genes (related to Fig. 2)**

**a** Cumulative PI plot comparing control and SMC1-depleted cells for upDEGs, nonDEGs, and downDEGs. *P* values were calculated using the two-sided Wilcoxon rank-sum test.

**b** Confirmation of SMC1 depletion by immunoblotting with polyclonal antibody against SMC1. The band intensity of SMC1 normalized to Ponceau was reduced to 8% after auxin treatment for 3 hr.

**c** Histograms of DNA content. Auxin treatment for 3 h to deplete SMC1 did not affect the cell cycle progression.

**d** Cumulative PI plot comparing control and SMC1-depleted cells for PI1-PI4 genes. *P* values were calculated using the two-sided Wilcoxon rank-sum test.

**e** Box plot showing the distribution of log2 fold change in PI upon SMC1 depletion at PI1-PI4 genes. A multiple Welch t-test was performed between every pair of gene groups. $^{**}P < 0.01$, $^{***}P < 0.001$. $P$ = 9.2e-5 (PI1 vs PI2), $P$ = 1.2e-7 (PI1 vs PI3), $P$ = 1.8e-10 (PI1 vs PI4), $P$ = 4.5e-3 (PI2 vs PI3), $P$ = 2.3e-6 (PI2 vs PI4).

**f** Comparison of PI and log2 fold change of PI in auxin-treated cells relative to control cells and distribution of the log2 fold change. *P* value and Spearman correlation coefficient (rho) were calculated using the Spearman rank correlation test. The 1112 and 3814 genes exhibited more than 1.1-fold increase and decrease in PI.

**g** Confirmation of Wapl KD by immunoblotting with polyclonal antibody against Wapl. The band intensity of Wapl normalized to Tubulin was reduced to 24% after siRNA treatment.

**h** Heatmaps showing signals of Rad21 in control and Wapl KD cells at 48999 Rad21 peaks sorted by Rad21 intensity.

**i** Box plot showing the distribution of log2 fold change in PI upon Wapl knockdown at PI1-PI4 genes. A multiple Welch t-test was performed between every pair of gene groups. *P* values less than 0.05 were not detected.

**j** Proportion of PI1-PI4 at PI-increased, PI-unchanged, and PI-decreased genes.

Fig. S3

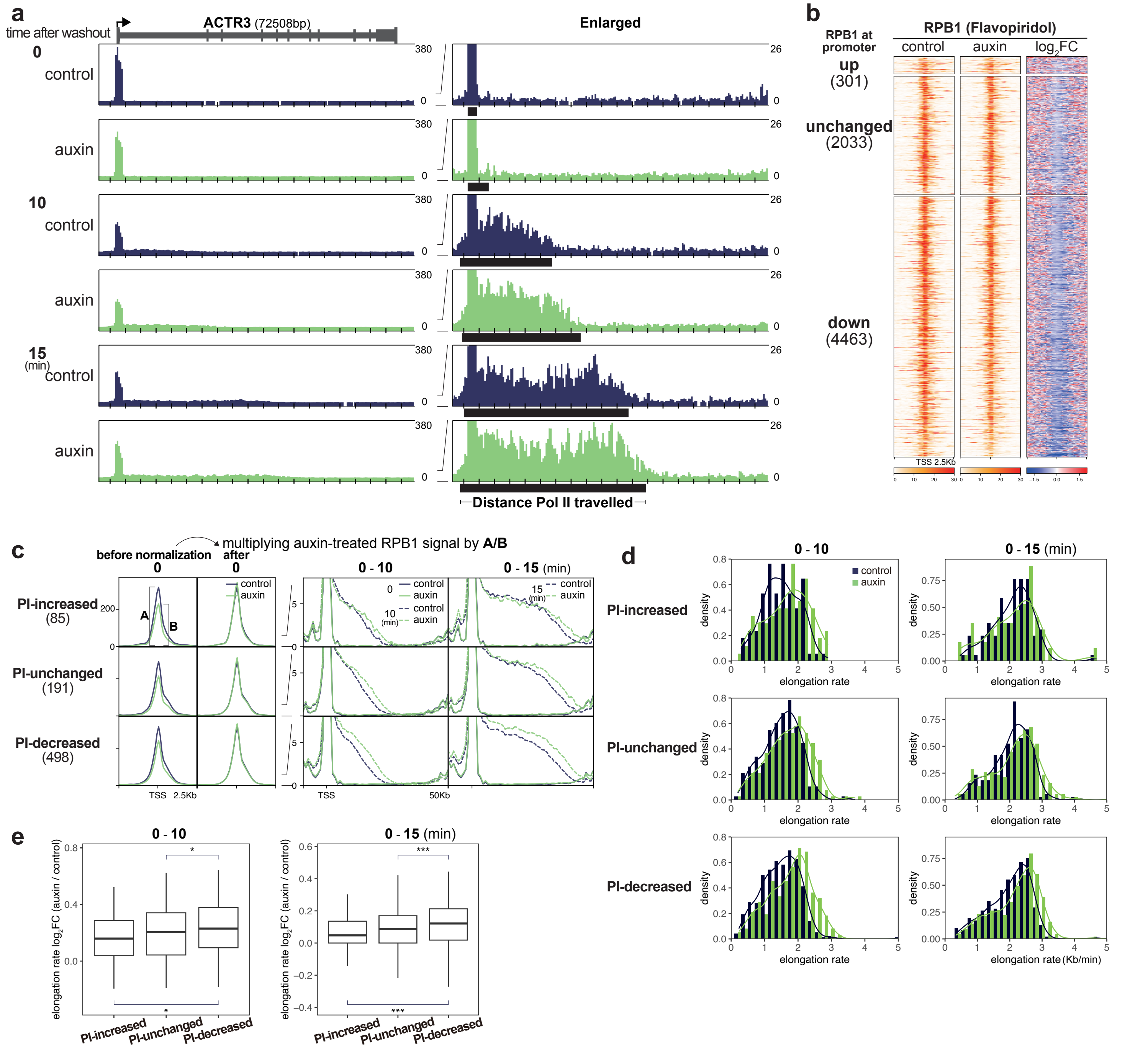


**Fig. S3: Cohesin restricts Pol II progression from promoters (related to Fig. 2)**

**a** Representative gene showing enhanced Pol II progression 10 and 15 min after washout of flavopiridol, an inhibitor of Cdk9. Black boxes indicate SICER-called peaks of RPB1, which correspond to the regions Pol II traveled.

**b** Heatmaps showing signals of RPB1 in flavopiridol-treated and flavopiridol plus auxin-treated cells at promoters of genes with up-regulated, unchanged, and down-regulated RPB1 promoter occupancy upon auxin treatment. Gene grouping and sorting orders are identical to those in Fig. 1b.

**c** Average plots of RPB1 0, 10, and 15 min after flavopiridol washout at PI-increased, PI-unchanged, and PI-decreased genes. Genes longer than 40kb and exhibiting SICER-called peaks of elongating RPB1 were analyzed. RPB1 signals in auxin-treated cells were multiplied by the ratio of flavppiridol-treated RPB1 intensity at each promoter in control cells to that in flavopiridol plus auxin-treated cells. Promoters were defined as regions within 1kb of the TSS of the genes. In the TSS-centered profile, the bin size was set to 100 bp, and in other profiles, the bin size was set to 1000 bp.

d Histograms showing elongation rates calculated based on the distances Pol II travelled at 10 and 15 min after flavopiridol washout at PI-increased, PI-unchanged, and PI-decreased genes.

e Box plot showing the distribution of log2 fold change in elongation rate upon auxin treatment at PI-increased, PI-unchanged, and PI-decreased genes. A multiple Welch t-test was performed between every two pair of gene groups. $*P < 0.05$, $***P < 0.001$. For 10 min, $P$ = 1.9e-2 (PI-increased vs PI-decreased), $P$ = 4.1e-2 (PI-unchanged vs PI-decreased). For 15 min, $P$ = 4.9e-4 (PI-increased vs PI-decreased), $P$ = 4.7e-4 (PI-unchanged vs PI-decreased).

### Fig. S4

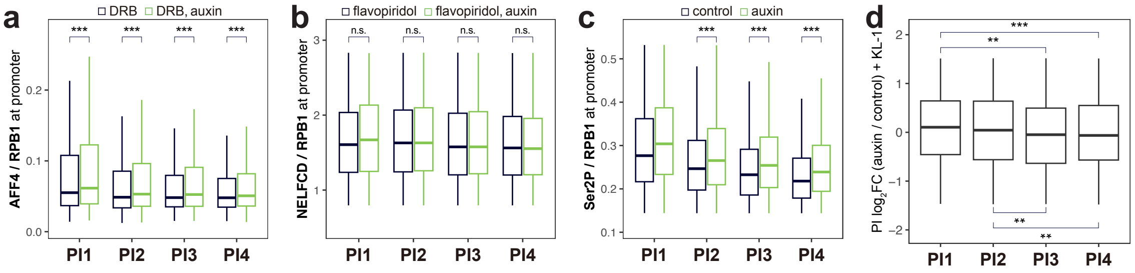


**Fig. S4: Cohesin restricts Pol II release from pausing by inhibiting SEC recruitment to promoters (related to Fig. 3)**

**a** Box plot showing the distribution of the relative amount of AFF4 to RPB1 at promoters (-500 to 500 bp) of PI1-PI4 genes in the presence of DRB. *P* values were calculated using the two-sided Wilcoxon rank-sum test between DRB-treated and DRB plus auxin-treated cells. ***$P < 0.001$. $P$ < 2.2e-16 (PI1), $P$ < 2.2e-16 (PI2), $P$ < 2.2e-16 (PI3), $P$ < 2.2e-16 (PI4).

**b** Box plot showing the distribution of the relative amount of NELFCD to RPB1 at promoters (-500 to 500 bp) of PI1-PI4 genes in the presence of flavopiridol. *P* values less than 0.05 were not detected using the two-sided Wilcoxon rank-sum test between flavopiridol-treated and flavopiridol plus auxin-treated cells.

**c** Box plot showing the distribution of log2 fold change in Ser2P levels upon auxin treatment at PI1-PI4 genes. *P* values were calculated using the two-sided Wilcoxon rank-sum test between control and auxin-treated cells. ***$P < 0.001$. $P$ < 2.2e-16 (PI2), $P$ < 2.2e-16 (PI3), $P$ < 2.2e-16 (PI4).

**d** Box plot showing the distribution of log2 fold change in PI upon auxin treatment in the presence of KL-1 at PI1-PI4 genes. A multiple Welch t-test was performed between every pair of gene groups. **$P < 0.01$, ***$P < 0.001$. $P$ = 1.3e-3 (PI1 vs PI3), $P$ = 3.1e-4 (PI1 vs PI4), $P$ = 8.6e-3 (PI2 vs PI3), $P$ = 1.1e-3 (PI2 vs PI4).

### Fig. S5

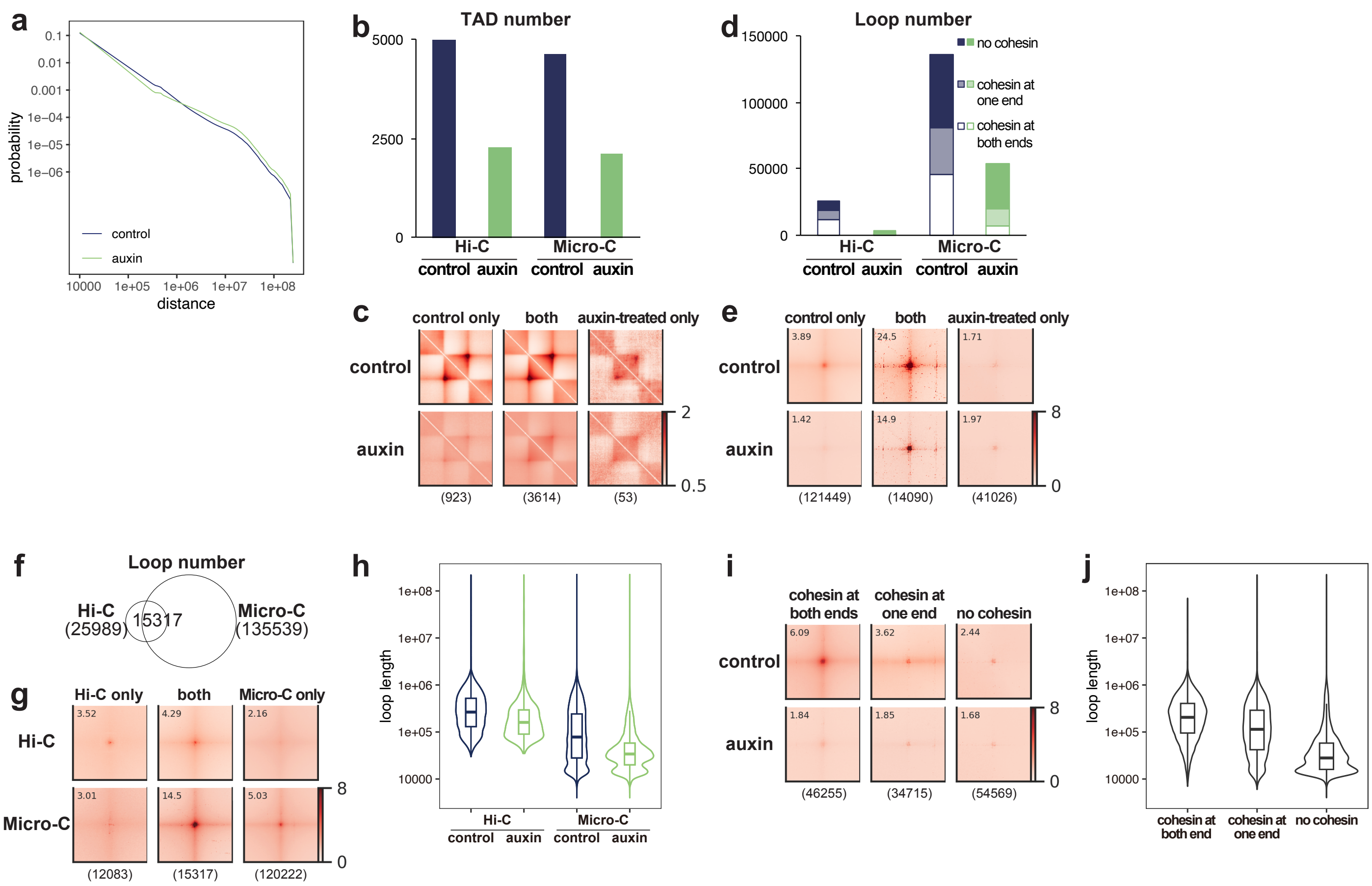


**Fig. S5: Micro-C analysis (related to Fig. 4)**

**a** Contact probability curves of control and auxin-treated cells.

**b** Total number of TADs detected by Hi-C and Micro-C in control and auxin-treated cells. The TADs were identified at 25kb resolution.

**c** Aggregate plots and numbers of Micro-C TADs shared or unique in control and auxin-treated cells.

**d** Total number of loops detected by Hi-C and Micro-C in control and auxin-treated cells. Hi-C loops were called at 5 kb resolution and Micro-C loops were called at 1k, 2k, and 5kb resolutions and merged.

**e** Aggregate plots of Micro-C loops shared or unique in control and auxin-treated cells.

**f** Venn diagram showing overlaps between Hi-C and Micro-C loops in control cells. Hi-C loops were called at 5 kb resolution and Micro-C loops were called at 1k, 2k, and 5kb resolutions and combined.

**g** Aggregate plots of loops in control cells shared and unique in Hi-C and Micro-C.

**h** Violin plot showing the distribution of loop lengths detected from Hi-C and Micro-C in control and auxin-cells.

**i** Aggregate plots of Micro-C loops with one, two, or no cohesin anchors.

**j** Violin plot showing the distribution of Micro-C loop lengths with one, two, or no cohesin anchors.

#### Fig. S6

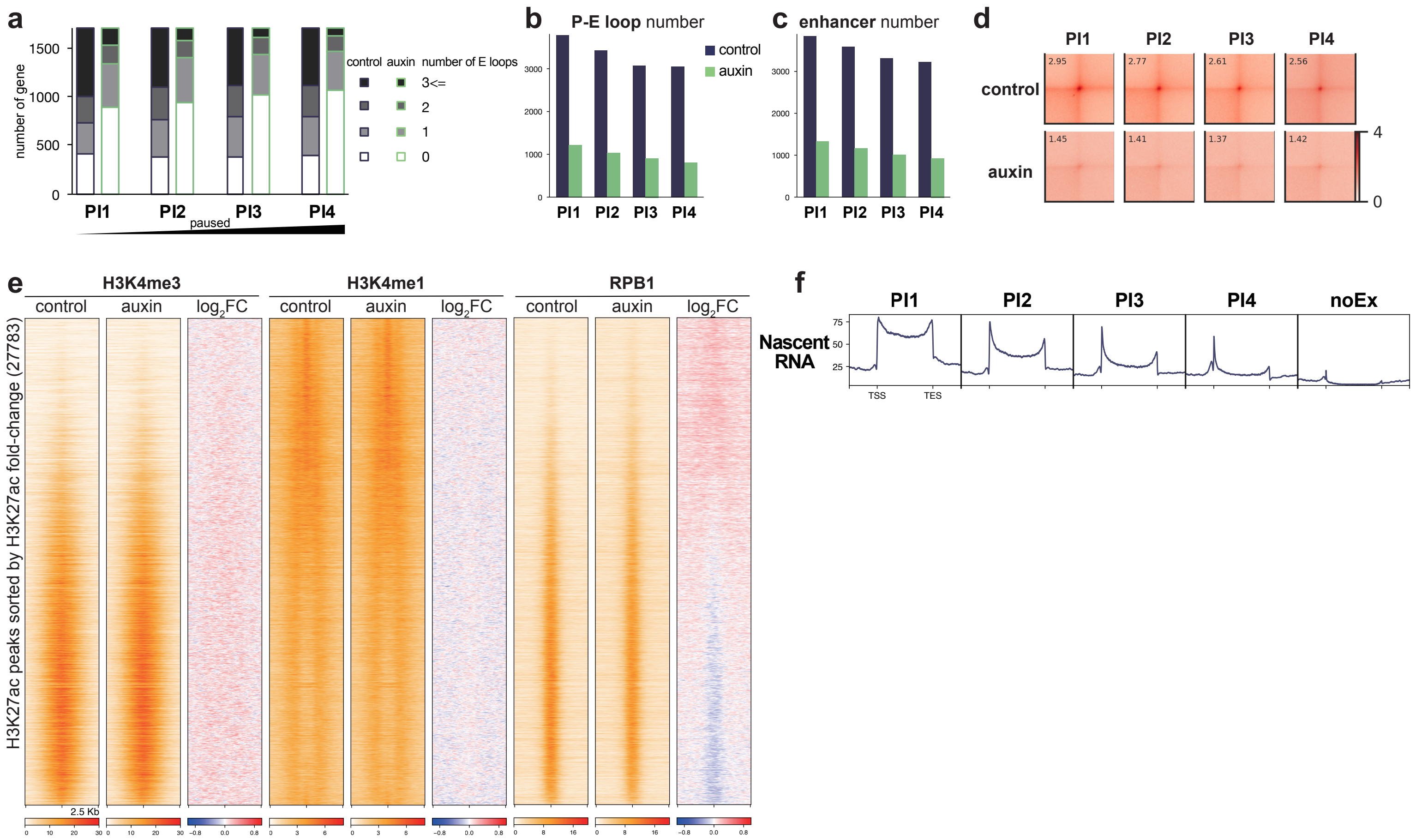


**Fig. S6: Lowly paused genes exhibited intensive contacts with multiple H3K27ac-enriched enhancers (related to Fig. 4)**

**a** Number of genes whose promoters were connected to enhancers via Micro-C loops identified in control and auxin-treated cells at PI1-PI4.

**b** Number of P-E (promoter-enhancer) loops anchoring at PI1-PI4 genes in control and auxin-treated cells.

**c** Number of enhancers connected to PI1-PI4 genes via P-E loops in control and auxin-treated cells.

**d** Aggregate plots of Micro-C loops between promoters of PI1-PI4 genes and enhancers in control and auxin-treated cells.

**e** Heatmaps showing signals of H3K4me3, H3K4me1, and RPB1 in control and auxin-treated cells at 27783 H3K27ac peaks. The sorting order is identical to that in Fig. 4d.

**f** Metagene plots of nascent RNA at PI1-PI4 and noEx. noEx were 1026 genes without H3K4me3 peaks within 5kb from TSSs nor any transcripts detected.

### Fig. S7

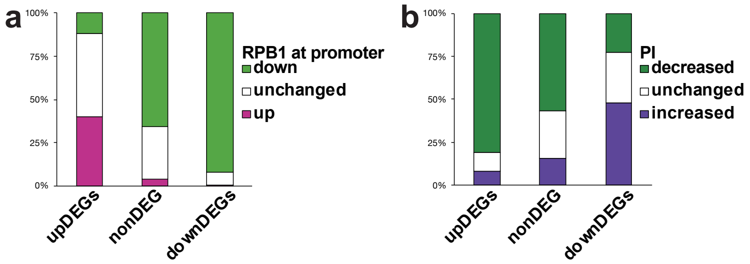


c

all gene

| PI \ RPB1 at promoter | up | - | down |
|---|---|---|---|
| increase | 1 | 4 | 11 |
| - | 1 | 7 | 19 |
| decrease | 3 | 18 | 35 |

**Fig. S7: Cohesin facilitates both Pol II binding at promoters and pausing simultaneously (related to Fig. 5)**

**a** The percentage of upDEGs, nonDEGs, and downDEGs at whose promoters RPB1 was increased or decreased after cohesin depletion.

**b** The percentage of PI-increased, PI-unchanged, and PI-decreased genes in upDEGs, unchanged genes, and downDEGs.

**c** Color scale table showing the proportions of the genes whose RPB1 amounts at promoters and PI were affected by cohesin depletion. The percentages were calculated for all genes analyzed in this study.
